# ILSM: A package for analyzing the interconnection structure of tripartite interaction networks

**DOI:** 10.1101/2024.11.18.623932

**Authors:** Weicheng Sun, Yangyang Zhao, Chuan Yan

**Affiliations:** Lanzhou University

**Keywords:** hybrid network, tripartite network, motif, interconnection pattern, R package, dual interaction, multiple interaction

## Abstract

1. In natural or human-disturbed ecosystems, ecological networks often comprise multiple interaction types, which have been often represented by multipartite ecological networks. One important aspect of their network architecture is how different interaction types or subnetworks are interconnected by shared species, here defined as the interconnection structure. Previous studies have proposed various indices of shared species to characterize macro-scale interconnection patterns and micro-scale centrality, but the meso-scale interconnection structures (here defined as interconnection motifs) remain largely unexplored. Furthermore, there is no package available in the R programming language for conducting analyses of various interconnection structures.
2. Within a tripartite network with two interaction subnetworks, we define the forms of interconnection motifs and unique roles within these motifs. Then we introduce the R package ILSM for analyzing interconnection patterns, interconnection centrality, and interconnection motif. Specifically, we derive mathematical expressions for the frequencies of interconnection motifs and species roles within motifs.
3. We describe the main functions in the package and demonstrate their uses with an example pollinator-plant-herbivore network.
4. ILSM will help ecologists understand how different types of interactions are interconnected together by shared species using interconnection pattern, centrality and motif.

## Background

Recently, ecological hybrid networks characterized by multiple interaction types have been a subject of great interest in ecology (Kéfi et al. 2012; Pocock et al. 2012a; Domínguez-García & Kéfi 2024). Many of these networks consist of several paired sets of species that form different bipartite networks, which are interconnected through shared species. These bipartite networks embedded in the hybrid networks can be represented by subnetworks (Fig. 1a)(Fontaine et al. 2011; Yan 2022). The subnetworks are often defined by trophic or mutualistic interactions with different functions; for instance, plant-pollinator and plant-herbivore subnetworks are interconnected by plants, or plant-insect and insect-parasitoid subnetworks are interconnected by herbivory insects (Domínguez-García & Kéfi 2024). Previous studies have analyzed the structures of tripartite networks with two interaction subnetworks, which offered novel insights into the complexity, functions and stability of natural communities (Sauve et al. 2016; Hervías-Parejo et al. 2020; Yan 2022). One important aspect in elucidating the architecture of these networks is how different interaction subnetworks are interconnected by connector species, which can be examined at macro-, micro-, and meso-scales (Fig. 1b-d).

**Fig. 1.**
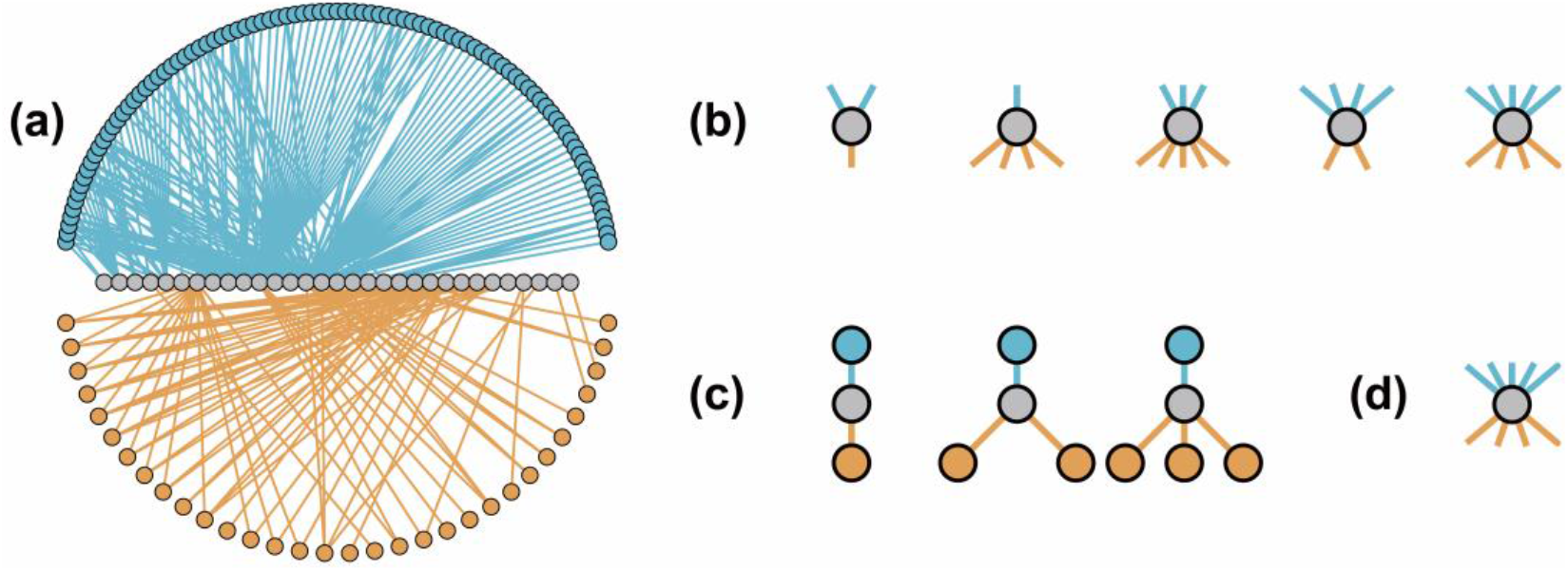
The visualization of an example tripartite interaction network (a, with three groups of species and two interaction subnetworks) and interconnection structures for connector species (b, macro-scale interconnection pattern; c, meso-scale interconnection motif; d, micro-scale interconnection centrality). Different colors of nodes indicate different groups of species. In panel a, the connector species have links from both subnetworks.

At the macro-scale, previous studies have proposed various interconnection patterns to describe the network-level interconnection structure of how shared species interconnect species from two different subnetworks in tripartite networks. (1) One basic feature is the proportion of connector species (species that have links in both subnetworks) in the shared set of species. For example, within a pollinator-plant-herbivore network where the plants interconnect pollinators and herbivores, not all plants may interact simultaneously with pollinators and herbivores. In the most extreme case, a value of 1 means all plants are involved in both subnetworks, and 0 means the two subnetworks are independent. (2) For the connector species, Sauve et al. (2016) employed two metrics for the interconnection patterns of networks: the correlation between connector species’ generalism in two subnetworks, and the correlation between the similarities of interacting species in two subnetworks among pairs of connector species. (3) The participation ratio refers to the average bias of interaction degree for connector species by quantifying whether the links of a node are primarily concentrated in one interaction subnetwork or if they are well distributed among the two subnetworks (Guimera & Amaral 2005; Battiston et al. 2014). (4) Domínguez-García and Kéfi (2024) proposed the proportion of shared species hubs (i.e. 20% of the shared species with the most connections) that are connectors nodes. At the micro-scale, centrality serves as a single measure of a node’s importance and has been frequently used to describe species importance and functions in ecological networks (Cagua et al. 2019; Mello et al. 2019).

The meso-scale structure in complex networks usually refers to motifs, which are simple subnetworks comprising a limited number of nodes. Motif analysis mainly explores the network properties by calculating the frequency of each motif and also counting the frequency with which nodes occur in different positions within motifs to quantify the species’ role in a network (Rodríguez-Rodríguez et al. 2017). Network motifs have been used to understand the functional properties of ecological networks, ecological processes and ecosystem stability (Stouffer & Bascompte 2010; Klaise & Johnson 2017; Simmons et al. 2021; Cirtwill & Wootton 2022). While the motifs have attracted significant interest and can be readily analyzed in ecological bipartite networks using the R software (Simmons et al. 2019b), this meso-scale interconnection structure within tripartite networks has been largely overlooked.

Recently, a study dissected pollination-predation hybrid networks into motifs with two or three species and found their motif profiles varied across agricultural management regimes (Martínez-Núñez & Rey 2021). Due to the limitation of species number in motifs, this work found only one pollination-plus-predation motif that involved two interaction types, however, a tripartite network might have various motifs interconnecting two subnetworks. For clarification, we define the interconnection motifs as the subgraphs that interconnect two interaction subnetworks, including species from both two subnetworks and connector species (Fig. 1c). A tripartite interaction network may exhibit a profile of interconnection motifs for each pair of connected subnetworks, i.e., the frequency of interconnection motifs. Then the role of each connector species can also be defined by counting the frequency with which species occur in various positions within interconnection motifs. To our knowledge, the different forms of interconnection motifs remain undefined in tripartite interaction networks.

Currently, there exist at least two significant gaps in the interconnection structures of tripartite interaction networks. First, in contrast to the various metrics on interconnection structures at micro- and macro scales, meso-scale interconnection structures have been largely unexplored. Second, no special software exists for analyzing various interconnection structures in ecological networks. In this study, we aim to fill the two gaps by defining interconnection motifs and introducing the R package ILSM that would facilitate the calculation of various metrics of interconnection structures in tripartite networks with two interaction types. First, we focus on the introduction and definition of interconnection motifs because the centrality and interconnection patterns have been previously established in the literature. Second, we provide a detailed description of primary functions to calculate the metrics. Third, we utilize our functions to analyze an empirical tripartite network. We note that the R package we have developed is tailored for tripartite ecological networks, which are characterized by two interaction types or subnetworks. However, it can apply to networks comprising four or more sets of species and subnetworks interconnected by shared species, as the networks with more subnetworks can be disaggregated into multiple tripartite networks. Different types of interactions can be represented by interaction layers (Pilosof et al. 2017; Hervías-Parejo et al. 2020).

However, the tripartite network in this study has no interlayer links, so our framework of analysis on interconnection structures is different from that of multilayer networks. Actually, plenty of multipartite networks with multiple interaction types have no interlayer links in previous studies (Pocock et al. 2012b; Domínguez-García & Kéfi 2024), and it is thus important to analyze how connector species interconnect different types of interactions in such networks. For clarification, we define the tripartite network as this: it is composed of three sets of nodes (*a*-nodes, *b*-nodes, and *c*-nodes) and two subnetworks (the *P* subnetwork with links between *a-*nodes and *b*-nodes, and the *Q* subnetwork with links between *b*-nodes and *c-*nodes); the *b*-nodes are the shared set of nodes; connector nodes are the common nodes of both subnetworks in the *b*-nodes (Fig. 1a).

## Methods and features

### Definition of interconnection motif

We define that an interconnection motif must comprise three sets of connected nodes: the connector nodes (belonging to *b*-nodes), the nodes in one subnetwork (belonging to *a*-nodes in the P subnetwork), and the nodes in the other subnetwork (belonging to *c*-nodes in the Q subnetwork). We further restricted each motif to contain six nodes, resulting in a total of 48 distinct motifs (Fig. 2). Permitting more than six nodes yields redundant motif forms that can be represented by aggregating motifs with six or fewer nodes. Variations in nodes and links across motifs alter the roles of connector nodes. Consequently, the connector nodes within the 48 motifs can be classified into 70 unique roles based on their interaction patterns with other nodes within the motifs. Nodes in a motif share the same position if a permutation of these nodes and their links does not modify the motif structure.

**Fig. 2.**
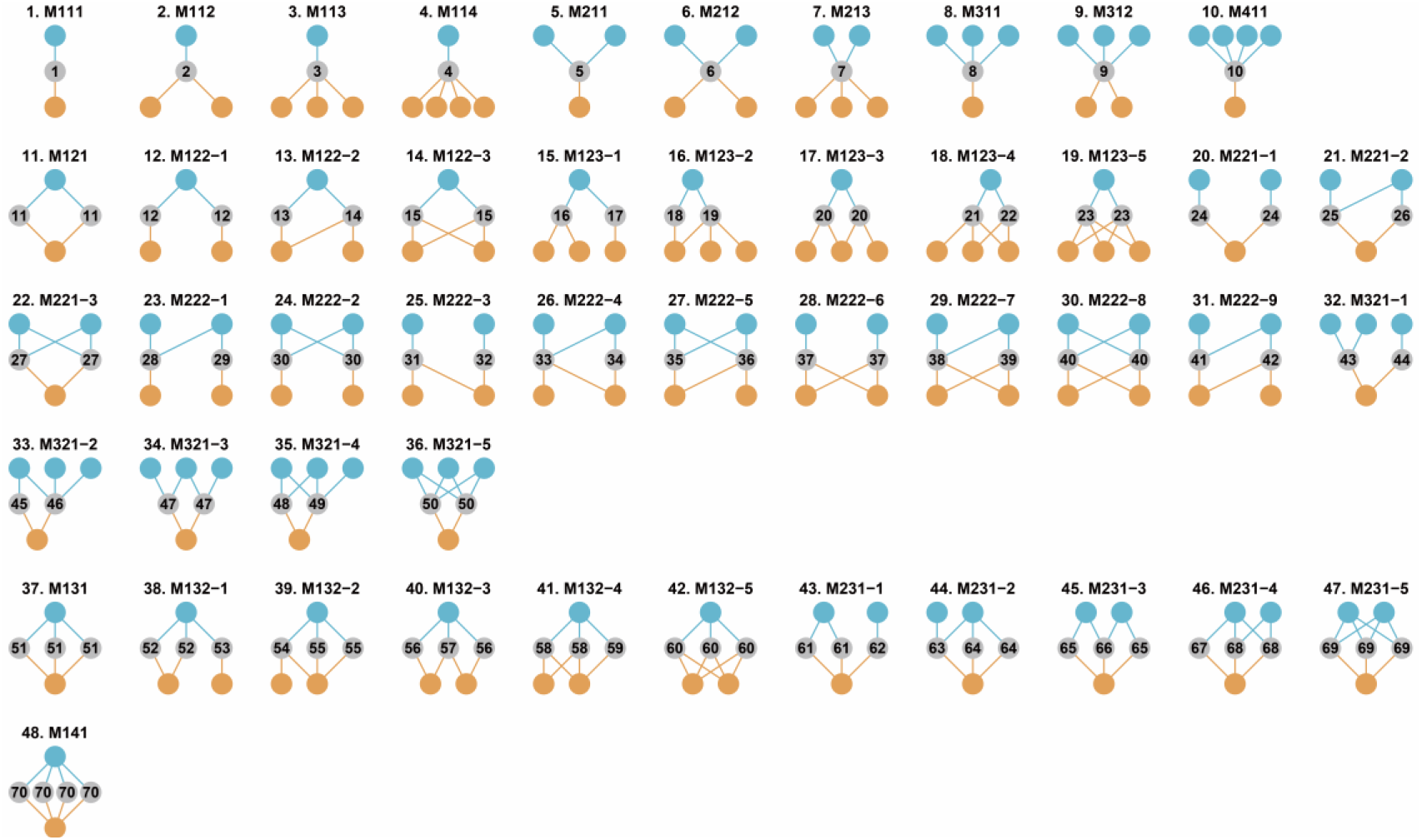
The 48 forms of interconnection motifs with 3-6 nodes. Blue and grey nodes form one subnetwork, and grey and orange nodes form the other subnetwork. Grey nodes are connector nodes and the dashed lines indicating the connector nodes in Fig. 1 were dropped for simplification. The motifs are named “MABC-i”: M means “motif’, “A” is the number of *a*-nodes, “B” is the number of *b*-nodes, “C” is the number of *c*-nodes and “i” is the serial number for the motifs with the same “ABC”. The interconnection motifs are ordered by the number of connector nodes (from 1 to 4). The numbers from 1 to 70 in connector nodes represent the unique roles.

Motifs can uncover indirect interactions present in the meso-scale topology of networks (Simmons et al. 2019a). In food webs or mutualistic networks, exploitative competition and apparent competition are typical indirect relationships revealed by three-species motifs. The interconnection motifs extend that to a tripartite version. For example, M212 represents an apparent-competition-plus-explorative-competition in a plant-herbivore-parasitoid network or facilitative-interaction-plus-explorative-competition in a pollinator-plant-herbivore network. We have developed two functions to quantify the frequency of different interconnection motifs and that of connector species’ motif roles.

### Functions for interconnection motifs

#### icmotif_count

The *icmotif_count* function calculates the frequency of each defined interconnection motif in a tripartite network. The function takes a matrix or graph data type (Appendix S1). The algorithm for counting interconnection motifs is designed by extending the fast approach from Simmons et al. (2019b), which uses mathematical operations directly on the bi-adjacency matrix. We drive 48 expressions to enumerate the frequencies of 48 interconnection motifs and explain them with examples in Appendix S2.

#### icmotif_role

The *icmotif_role* function calculates the frequency of each role within interconnection motifs for each connector species using a similar approach as above. We also drive 70 expressions to enumerate the frequencies of 70 roles and explain them with examples in Appendix S3.

### Functions for interconnection patterns

#### poc

The *poc* function calculates the proportion of connector nodes in the shared set of nodes (POC).

#### coid and cois

The *coid* function calculates the correlation of interaction degree (CoID) and the *cois* function calculates the correlation of interaction similarity (CoIS) for connector species. For CoID, two types of degrees (*d*_*P*_ from P and *d*_*Q*_ from Q) are calculated for each connector species, and the correlation of two degrees is calculated. For CoIS, the pairwise Jaccard index of interaction partners for shared species in each subnetwork is measured to form two dissimilarity matrices, and then the Mantel correlation coefficient between them is calculated.

#### pc

The *pc* function calculates the participation coefficient. For each connector node *i, pc*_*i*_ is calculated as two times the ratio between the lowest degree in both interaction subnetworks (d_P_ and d_Q_) divided by the total degree of node *i* :

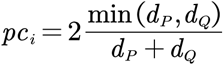

Hence, the participation coefficient for all connector nodes (PCc) is represented by the average value of all *pc*_*i*_.

#### hc

The *hc* function counts the proportion of connector nodes in connector node hubs (HC). The connector node hubs are the top x% of shared nodes with the highest degree (Domínguez-García and Kéfi 2021). The default x% is 20%.

### Functions for interconnection centrality

The *node_icc* function measured seven centrality of connector nodes, so-called interconnection centrality. This function is simply developed following the R package *igraph*, with seven centrality measures that have been commonly applied to networks, i.e., degree, PageRank, hub, authority, betweenness, eigenvector, and closeness centrality (Page et al. 1999; Magnani et al. 2013; De Domenico et al. 2015).

### Example

#### A worked example

As a worked example, we used a published pollinator-plant-herbivore (PPH) tripartite network (Villa-Galaviz et al. 2021). This PPH network is a subset of a large hybrid network including plants, flower visitors, leaf minors, and parasitoids from a long-term nutrient manipulation experiment (Colt Park Meadows) located at 300 m elevation in the Ingleborough National Nature Reserve in the Yorkshire Dales, northern England (54°12′N, 2°21′W). It contained 93 species of pollinators, 31 species of plants and 32 species of herbivores, corresponding with 188 mutualistic interactions in the pollinator-plant subnetwork and 80 antagonistic interactions in the plant-herbivore interactions.

Here we used the ILSM package to analyze and visualize interconnection structures of this PPH network. All five interconnection patterns were shown in Fig. 3a (POC = 0.26, CoID = 0.17, CoIS = 0.48, PCc = 0.38, HC = 0.67). The low POC indicates only a small proportion of shared species have interactions in both subnetworks (i.e., connector species) and for connector species, the correlation of their degrees and interaction similarity from the two subnetworks are both positive but low. The high HC indicates connector nodes are mostly connection hubs in the network. The centrality analysis reveals a spectrum of values across different indices and among connector species (Fig. 3b). Three connector species (*Ranunculus acris, Leucanthemum vulgare*, and *R. repens*) show high levels in all seven centrality metrics, and the rest exhibit low centralities except for closeness centrality.

**Fig. 3.**
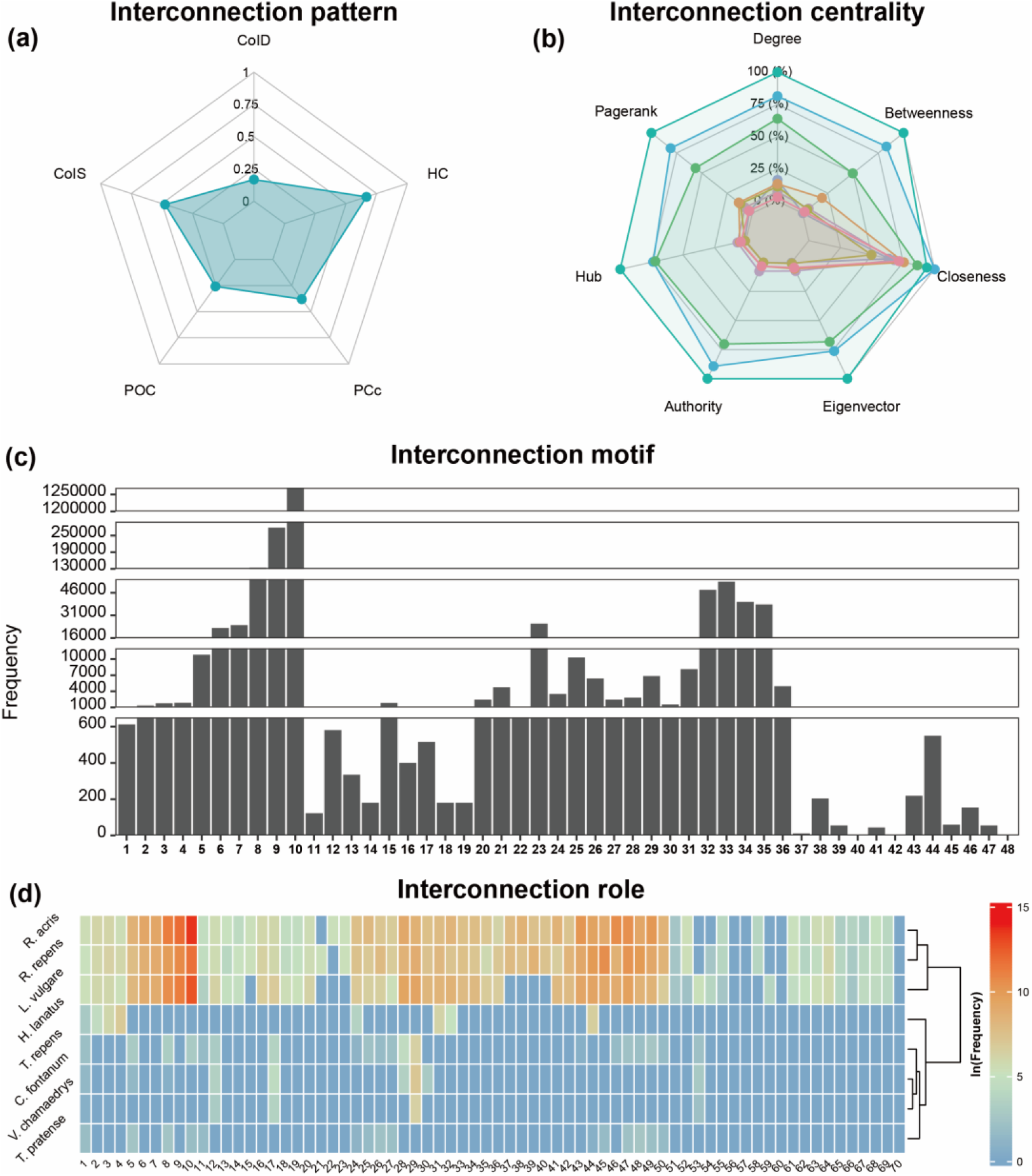
The interconnection structures of the example PPH network. (a) Five interconnection patterns. (b) Seven centrality indices for eight connector species. (c) The frequencies of 48 interconnector motifs. (d) The frequencies (ln-transformed) of 70 roles for eight connector species in the interconnection motifs.

For interconnection motifs, the frequencies vary substantially across 48 motifs, and those with the most connector species are rarely detected (Table S1, Fig. 3c). The 10th motif “M411” has the highest number probably because of the vast possibilities of connecting pollinators with plants. The role profiles of connector species indicate a few species have much more frequent roles within interconnection motifs than the rest, and species can be grouped based on cluster analysis of roles (Fig. 3d, Table S2).

#### Implementation and availability

The ILSM package is available for the R programming language. The installation, manual and source code of the package can be achieved from the Github page: https://github.com/WeichengSun/ILSM.

## Conclusion

ILSM is an R package for analyzing the interconnection structure of shared species in tripartite networks. On the whole, this package collates some of the computational measures for interconnection patterns and centrality indices and also defines interconnection motifs with functions to analyze them. At the network and species levels, interconnection motifs can reflect extra differences beyond interconnection patterns and centrality indices in tripartite networks with two interaction types or subnetworks. There might be two relevant applications in future studies. First, the interconnection patterns and motif profiles might be used to investigate the biotic and abiotic drivers for the variation of the architecture of tripartite interaction networks on various environmental gradients. Second, within a network, the interconnection centrality and motif roles of species may help to understand the ecological and evolutionary factors affecting species’ importance and function across different interaction subnetworks. Upon the recent surge of studies in hybrid ecological networks, we hope this package will assist with related research and inspire new perspectives.

## References

Battiston, F., Nicosia, V. & Latora, V. (2014) Structural measures for multiplex networks. Physical Review E, 89, 032804.

Cagua, E.F., Wootton, K.L. & Stouffer, D.B. (2019) Keystoneness, centrality, and the structural controllability of ecological networks. Journal of Ecology, 107, 1779–1790.

Cirtwill, A.R. & Wootton, K. (2022) Stable motifs delay species loss in simulated food webs. Oikos, 2022.

De Domenico, M., Solé-Ribalta, A., Omodei, E., Gómez, S. & Arenas, A. (2015) Ranking in interconnected multilayer networks reveals versatile nodes. Nature Communications, 6, 6868.

Domínguez-García, V. & Kéfi, S. (2024) The structure and robustness of ecological networks with two interaction types. PLOS Computational Biology, 20, e1011770.

Fontaine, C., Guimarães Jr, P.R., Kéfi, S., Loeuille, N., Memmott, J., van Der Putten, W.H., Van Veen, F.J. & Thébault, E. (2011) The ecological and evolutionary implications of merging different types of networks. Ecology Letters, 14, 1170–1181.

Guimera, R. & Amaral, L.A.N. (2005) Cartography of complex networks: modules and universal roles. Journal of Statistical Mechanics: Theory and Experiment, 2005, P02001.

Hervías-Parejo, S., Tur, C., Heleno, R., Nogales, M., Timóteo, S. & Traveset, A. (2020) Species functional traits and abundance as drivers of multiplex ecological networks: first empirical quantification of inter-layer edge weights. Proceedings of the Royal Society B-Biological Sciences, 287.

Kéfi, S., Berlow, E.L., Wieters, E.A., Navarrete, S.A., Petchey, O.L., Wood, S.A., Boit, A., Joppa, L.N., Lafferty, K.D. & Williams, R.J. (2012) More than a meal… integrating non-feeding interactions into food webs. Ecology letters, 15, 291–300.

Klaise, J. & Johnson, S. (2017) The origin of motif families in food webs. Scientific Reports, 7, 16197.

Magnani, M., Micenková, B. & Rossi, L. (2013) Combinatorial analysis of multiple networks. ArXiv, abs/1303.4986.

Martínez-Núñez, C. & Rey, P.J. (2021) Hybrid networks reveal contrasting effects of agricultural intensification on antagonistic and mutualistic motifs. Functional Ecology, 35, 1341–1352.

Mello, M.A., Felix, G.M., Pinheiro, R.B., Muylaert, R.L., Geiselman, C., Santana, S.E., Tschapka, M., Lotfi, N., Rodrigues, F.A. & Stevens, R.D. (2019) Insights into the assembly rules of a continent-wide multilayer network. Nature Ecology and Evolution, 3, 1525–1532.

Page, L., Brin, S., Motwani, R. & Winograd, T. (1999) The PageRank Citation Ranking : Bringing Order to the Web. The Web Conference.

Pilosof, S., Porter, M.A., Pascual, M. & Kefi, S. (2017) The multilayer nature of ecological networks. Nature Ecology and Evolution, 1, 101.

Pocock, M.J., Evans, D.M. & Memmott, J. (2012a) The robustness and restoration of a network of ecological networks. Science, 335, 973–977.

Pocock, M.J., Evans, D.M. & Memmott, J. (2012b) The robustness and restoration of a network of ecological networks. Science, 335, 973–977.

Rodríguez-Rodríguez, M.C., Jordano, P. & Valido, A. (2017) Functional consequences of plant-animal interactions along the mutualism-antagonism gradient. Ecology, 98, 1266–1276.

Sauve, A.M.C., Thebault, E., Pocock, M.J.O. & Fontaine, C. (2016) How plants connect pollination and herbivory networks and their contribution to community stability. Ecology, 97, 908–917.

Simmons, B.I., Beckerman, A.P., Hansen, K., Maruyama, P.K., Televantos, C., Vizentin-Bugoni, J. & Dalsgaard, B. (2021) Niche and neutral processes leave distinct structural imprints on indirect interactions in mutualistic networks. Functional Ecology, 35, 753–763.

Simmons, B.I., Cirtwill, A.R., Baker, N.J., Wauchope, H.S., Dicks, L.V., Stouffer, D.B. & Sutherland, W.J. (2019a) Motifs in bipartite ecological networks: uncovering indirect interactions. Oikos, 128, 154–170.

Simmons, B.I., Sweering, M.J.M., Schillinger, M., Dicks, L.V., Sutherland, W.J. & Di Clemente, R. (2019b) bmotif: A package for motif analyses of bipartite networks. Methods in Ecology and Evolution, 10, 695–701.

Stouffer, D.B. & Bascompte, J. (2010) Understanding food-web persistence from local to global scales. Ecology letters, 13, 154–161.

Villa-Galaviz, E., Smart, S.M., Clare, E.L., Ward, S.E. & Memmott, J. (2021) Differential effects of fertilisers on pollination and parasitoid interaction networks. Journal of Animal Ecology, 90, 404–414.

Yan, C. (2022) Nestedness interacts with subnetwork structures and interconnection patterns to affect community dynamics in ecological multilayer networks. Journal of Animal Ecology, 91, 738–751.

